# Cryo-EM mapping of circumsporozoite protein epitope accessibility and visualization of antibody-induced shedding on *Plasmodium falciparum* sporozoites

**DOI:** 10.64898/2026.09.04.749346

**Authors:** Nancy Hom, Connor Weidle, Marie Pancera, Kelly K. Lee

**Affiliations:** Department of Medicinal Chemistry, University of Washington, Seattle, WA 98195; Vaccine and Infectious Disease Division, Fred Hutchinson Cancer Center, Seattle, WA 98109; Department of Biochemistry, University of Washington, WA 98195

## Abstract

The circumsporozoite protein (CSP) on the surface of the sporozoite stage of *Plasmodium falciparum* parasites is one of the major malaria antigens recognized by the adaptive immune system. Antibodies against CSP can provide protection from malaria infection following mosquito bite of the human host. Recent studies have shown how such antibodies recognize recombinant, truncated forms of *Plasmodium falciparum* CSP (PfCSP); however, much remains unknown about epitope accessibility and presentation of PfCSP on the parasite itself. Understanding the nature of the native antigen is of significant interest for optimization of immunogen design, since immunization with whole attenuated sporozoites has been shown to provide long-lasting protection against malaria disease. Here we use cryo-electron microscopy and tomography to investigate how antibodies against various PfCSP epitopes (N-terminal domain (NTD), Junction, central repeat, and C-terminal domain (CTD)) impact the PfCSP coat on whole *Plasmodium falciparum* sporozoites (PfSPZ). We observe that PfCSP by itself cannot be visualized on the surface of the cell by cryo-EM. However, by titrating in antibodies that recognize the Junction motif and central repeats, we delineate a boundary layer formed by PfCSP on the sporozoite outer membrane. The antibody concentration series also enabled us to determine conditions required to induce the dramatic CSP shedding reaction. By comparing matched monovalent FAb with IgG, we demonstrate that antibody bivalency is critical to driving this reaction. Additionally, we demonstrate that even at high concentrations, antibody against the NTD does not bind, indicating the absence or inaccessibility of this epitope on the PfCSP displayed by radiation-attenuated sporozoites, while the CTD can be bound by antibodies without producing a CSP shedding reaction. These studies shed light on the nature of the PfCSP coat in its native context on the parasite, thus revealing the sporozoite’s antigenic profile.

**AUTHOR SUMMARY:** Studies of anti-malaria antibodies targeting the circumsporozoite protein (CSP), the dominant antigen on the surface of sporozoites that are transmitted from mosquito to human hosts, have focused primarily on their interaction with recombinant CSP fragments. A number of such antibodies have been shown to provide protection in animal models and in early human clinical trials. However, little is known about how the antibodies directly interact with CSP on whole sporozoites (SPZ). Using cryo-electron microscopy and cryo-electron tomography we examined how antibodies against major CSP epitopes recognize native CSP on intact *Plasmodium falciparum* SPZ. Understanding the accessibility of CSP epitopes on SPZ can illuminate the nature of the native antigen and indicate which epitopes are most effective to target through vaccination in order to produce a specific functional response such as CSP shedding. Additionally, we investigated the effects of antibody concentration and bivalency on antibody-induced CSP crosslinking and shedding. These findings can guide future vaccine immunogen design and help identify mechanisms of action for CSP-targeting antibodies.

## INTRODUCTION

Malaria imposes a significant disease burden worldwide through infection of an estimated 249 million people, leading to over 600,000 deaths per year [1]. Caused by a mosquito-borne *Plasmodium* parasite, a major trait of these infections is that individuals can become re-infected despite developing antibodies and mounting robust humoral immune responses [2]. Though they are an important milestone towards developing improved efficacy and longevity of protection against malaria, recent vaccines using recombinant *Plasmodium* surface antigens have shown only partial protection through elicitation of antibodies without long-lasting durability [3–6]. By contrast, inoculation with whole, attenuated *Plasmodium* parasites has been shown to yield durable and more efficacious immune responses [7–12], possibly due to responses from tissue- resident T-cells in the liver [2, 13, 14]. Understanding the antibody response to malaria infection and to vaccine antigens is essential for developing more effective means of eliciting desirable, efficacious antibody responses with more potent vaccines. Additionally, it is necessary to better understand the effect that anti-malarial antibodies have on the parasite in order to identify their mechanisms of inhibition.

The sporozoite (SPZ) stage of the *Plasmodium* life cycle represents an opportune target for blocking by antibodies [15, 16]. SPZ form in the mosquito midgut then migrate to the insect’s salivary glands where they remain until they are delivered into the mammalian host by a mosquito bite. Once introduced into the host dermis, sporozoites migrate to the circulatory system, which delivers them to the liver where they traverse through several cells before subsequently invading a hepatocyte; then the plasmodium replicates asexually to produce merozoites that are released into the blood where red blood cells support latter stages of reproduction and infection of mosquito vectors [17, 18]. By blocking the parasite at the SPZ stage, infection can be prevented [10, 15, 19, 20].

Circumsporozoite protein (CSP) is the most abundant protein on the SPZ surface. It is small (44kDa) and expected to be largely disordered due to its abundance of proline-rich repeats. It is composed of three main domains: an NTD that has eluded structural characterization, a central repeat region comprised primarily of dozens of NANP serial repeat motifs interspersed with NVDP minor repeats and a unique NPDP sequence, and a CTD that partially adopts a globular alpha-TSR-like fold [21] and is glycosylphosphatidylinositol (GPI)-anchored at its C- terminus to the outer SPZ membrane [22] (**Fig 1A**). CSP is essential for midgut to salivary gland migration in the mosquito host, as well as intradermal motility in the mammalian host and hepatocyte invasion [17, 23]. The link between CSP domains, structures, and functions as the SPZ travels from the mosquito midgut to the mammalian liver remain areas of speculation. For example, it has been reported for CSP that the N-terminal domain (NTD) is needed to mask the C-terminal domain (CTD) before the SPZ reaches the liver as an immune evasion strategy, and that a proteolytic cleavage removes the NTD, allowing the CTD to interact with the heparin sulfate on the surface of hepatocytes for invasion and the progression to the next stages of the parasite lifecycle [24–34]. It is unclear, however, at which point this cleavage happens and what forms of CSP are present on whole sporozoites. Indeed, reports are in conflict regarding whether NTD-targeting antibodies bind to whole sporozoites [31, 34, 35]. A clearer understanding of which domains are accessible on the SPZ surface is needed to guide next-generation vaccine development.

**Fig 1.**
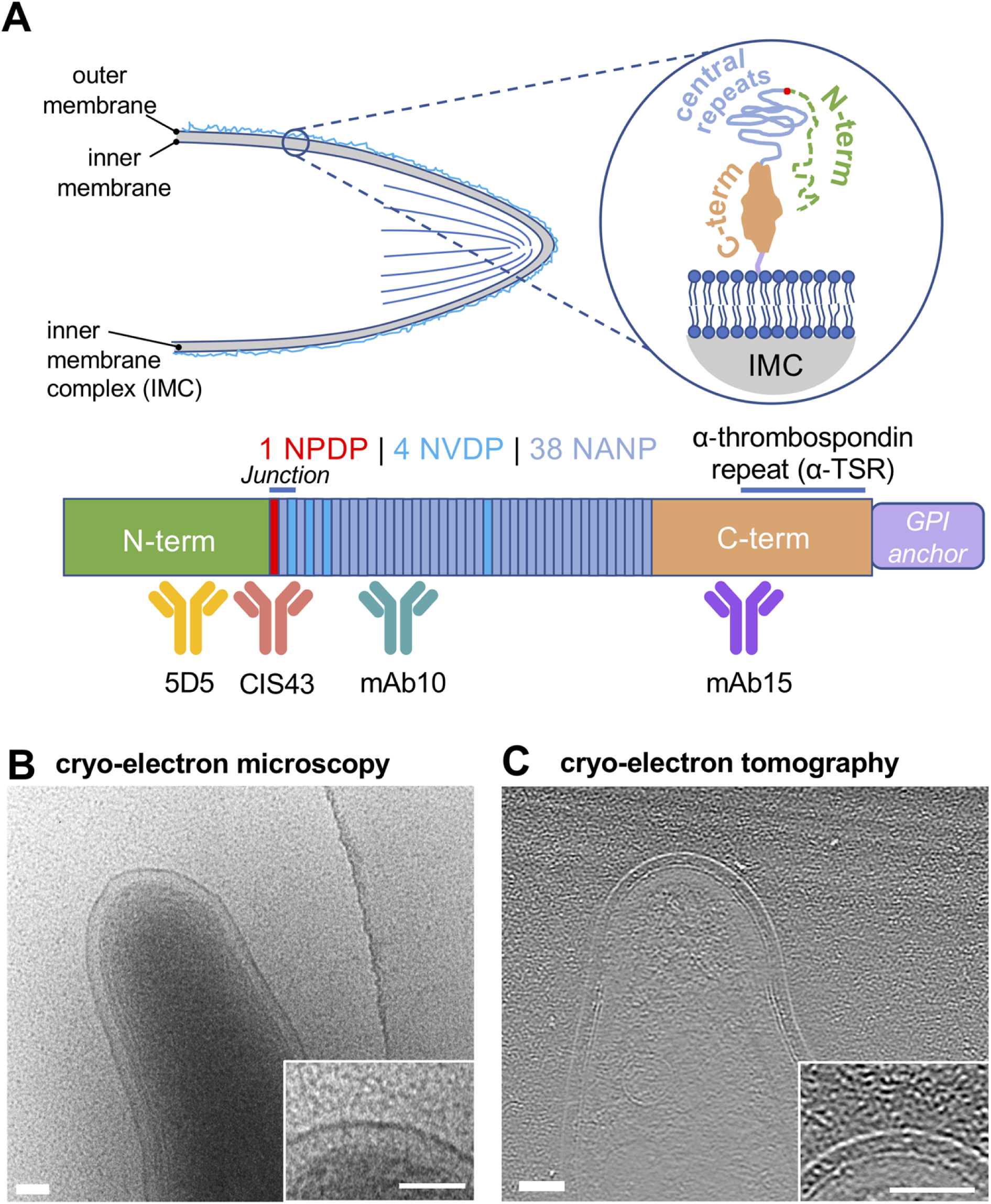
Antibody-naïve *Plasmodium falciparum* sporozoites are thought to be coated by the circumsporozoite protein (CSP). **(A)** Schematic of the PfSPZ surface with disordered PfCSP and linear depiction of PfCSP protein with NTD (green), central repeats (minor NVDP repeats and major NANP repeats, blue), CTD (orange), and GPI anchor (lavender). The PfCSP Junction is identified by the inclusion of the unique sequence NPDP (red). The CSP model is based on previous reports and the observations presented in this study. **(B)** Low-dose cryo-electron microscopy image of a radiation-attenduated *P. falciparum* sporozoite (PfSPZ); inset shows close up of PfSPZ surface with no PfCSP (44 Kda) visible, while human serum albumin (HSA, 60 kDa) are observed in the background as punctate granular features. **(C)** Cryo-electron tomography of a SPZ with the central tomographic slice shown; inverted contrast; inset of surface with no CSP visible, while HSA is observed as punctate granular features in the background. Additional views of PfSPZ with no antibody bound are presented in Supplemental Figure S1. All scalebars 100 nm.

It has long been recognized that CSP is the immunodominant antigenic protein on sporozoites, but only recently have structural studies revealed antibody-antigen recognition at high resolution and provided more detailed understanding of epitope specificity in the immune response [36–40]. While monoclonal antibodies that target each of the 3 domains have been isolated over the past several years, the central repeat region appears to be the most immunodominant [41–43]. The majority of the antibodies generated by natural infection, and the majority of those that are protective, bind to the proline-rich repeat motifs in the central segment of the CSP protein [23].

Antibodies against the NTD and CTD domains tend to be less abundant, which has led to speculation that these domains are not accessible or have limited binding in native CSP [23, 28, 31]. In the case of NTD-binding antibodies, it is unclear whether this epitope is available on SPZ and may be dependent on source of sporozoite (midgut or salivary glands) [31, 34–36, 44]. Indeed, studies diverge over whether antibodies that target epitopes within the NTD are able to bind to CSP on whole sporozoites and whether they are protective or not [30, 35]. Recent reports also have identified antibodies against the CSP Junction region and nearby minor NVDP repeats that separate the NTD and central repeat region. Some of these antibodies have shown promising protection profiles in human clinical trials (CIS43, L9) [36, 45–47], pointing to these epitopes as being highly desirable targets for vaccine targeting, though how this class of antibodies interact with whole sporozoites remain to be seen.

In most cases, CTD-binding antibodies appear to not provide protection from sporozoite infection alone [23, 48, 49], though recently antibodies against the CTD have been shown to be associated with some protection against malaria challenges in combination with Junction-targeting antibodies [50], and Beutler et al. reported a CTD-binding antibody that can inhibit malaria infection in a murine model [51]. Generally, our understanding of antibody recognition of the CSP CTD and NTD, particularly in their native context on sporozoites, remains poorly understood.

Although crystallography and single particle cryo-EM have revealed the detailed structures of several anti-repeat antibodies in complex with truncated recombinant *Plasmodium falciparum* CSP (PfCSP), the available structures have not provided information for how these prevent infection by the parasite [36, 37, 39, 40]. Two key missing pieces of analysis are demonstration of how the repeat antibodies recognize native PfCSP on the surface of intact sporozoites and the effects they produce, as well as determination of whether recombinant PfCSP recapitulates PfCSP on the surface of *Plasmodium falciparum* sporozoites (PfSPZ). On the surface of whole PfSPZ, antibodies targeting the central repeats have been shown, under certain conditions, to produce a plume of density reflecting a shedding of the CSP coat in a phenomenon called the CSP reaction [52]. It is thought this happens when bivalent IgG bind and crosslink to multiple copies of CSP on the surface of the SPZ [53]. Although the shedding in the CSP reaction is a known phenomenon, the concentration range of antibodies that elicit this response is not well understood, nor is it understood whether shedding is essential for antibody-mediated protection. In a study from Aliprandini et al investigating how *P. yoelii* CSP shedding is connected to SPZ death, shedding of CSP and death of the SPZ have been linked, though parasite death has been shown to also require cell-terminating effectors that form pores in the SPZ membranes [54].

Here, we sought to gain a better understanding of the nature of native PfCSP on the *Plasmodium falciparum* sporozoite surface, to determine which PfCSP domains are accessible on intact cells, and to image the impact of antibodies engaging with PfCSP on the cells. To this end, we have used cryo-electron microscopy (cryo-EM) and cryo-electron tomography (cryo-ET) to image live, radiation-attenuated PfSPZ cells [10, 13, 55] incubated with a panel of anti-CSP antibodies across a range of concentrations. The antibodies we examine target the different domains (**Fig 1**) of PfCSP and include protective as well as non-protective antibodies. We have also compared the effect of FAb vs IgG to assess the role of antibody bivalency on the CSP reaction. Together, these results shed light on the organization of native PfCSP on the surface of sporozoites and the disposition and accessibility of the three major PfCSP domains, while also revealing the impact of antibody binding to this major antigenic malaria determinant in the context of whole sporozoites.

## RESULTS

### Density attributable to PfCSP is not visible on the PfSPZ surface

We first used cryo-electron microscopy to image whole PfSPZ in the absence of antibodies. Under the imaging conditions of relatively high defocus and low dose transmission electron microscopy (TEM) we used, where one might anticipate observing density for a protein coat that could be attributable to PfCSP on the sporozoite surface, no clearly discernable features were observed (**Fig 1B, S1 Fig**) beyond the apparently smooth outer membrane. A relatively uniform “inner membrane complex” compartment could be clearly discerned between the inner and outer membrane of the cell (**Fig 1B, S1 Fig**). Internal organelles and densely packed vesicles corresponding to rhoptries and micronemes were also visible in some cases, but considerable variation was observed in internal features observed from cell to cell. Some of this variation is likely due to which end of a cell was being imaged. For example, structures such as the apical ring complex that organizes microtubules could be observed in a small subset of sporozoites, but not in all fields of view (**S2 Fig**).

We reasoned that cryo-ET imaging, which provides 3D imaging to complement low dose projection TEM imaging, potentially could improve visualization of any protein on the cell surface. However, as was seen by cryo-EM, cryo-ET also showed SPZ with a smooth outer membrane with no discernable surface protein density (**Fig 1C)**. PfCSP is approximately 44 kDa in molecular weight, and it has been estimated that there are 10^5^-10^6^ copies per cell [56, 57]. By comparison, 60 kDa human serum albumin that was not fully washed away during SPZ pelleting could be identified in the background as granular densities (**Fig 1B and 1C**). These data suggest that the PfCSP protein is either absent from the antibody-naïve sporozoite surface, or PfCSP is highly disordered, lacking in a well-formed globular structure. Indeed, a predominantly disordered structure for PfCSP is anticipated given the low-complexity nature of the central proline-rich repeat region and would be consistent with previous reports by atomic force microscopy and small-angle X-ray scattering [58].

### Anti-repeat antibodies trace the extent of the PfCSP layer on PfSPZ surface

To identify which epitopes on PfSPZ-displayed CSP are accessible to antibodies, we next incubated PfSPZ with monoclonal antibodies against the central repeat region, NTD, and CTD. Antibody concentrations ranging from 0.26 to 130 µg/mL were examined since this range spans plausible antigen-specific antibody concentrations that may be found in human sera [59–63].

We first examined two PfCSP central repeat-binding antibodies, CIS43 (**Fig 2**) and mAb10 IgG (**Fig 3**). CIS43 is reported to have a bispecific recognition, binding preferentially to an epitope containing NPDP and the NVDP minor repeat, which overlaps with the Junction region between the NTD and central repeat region, while its secondary binding targets the major NANP central repeat region; mAb10 by contrast is strictly selective for the NANP major repeat sequence [36].

**Fig 2.**
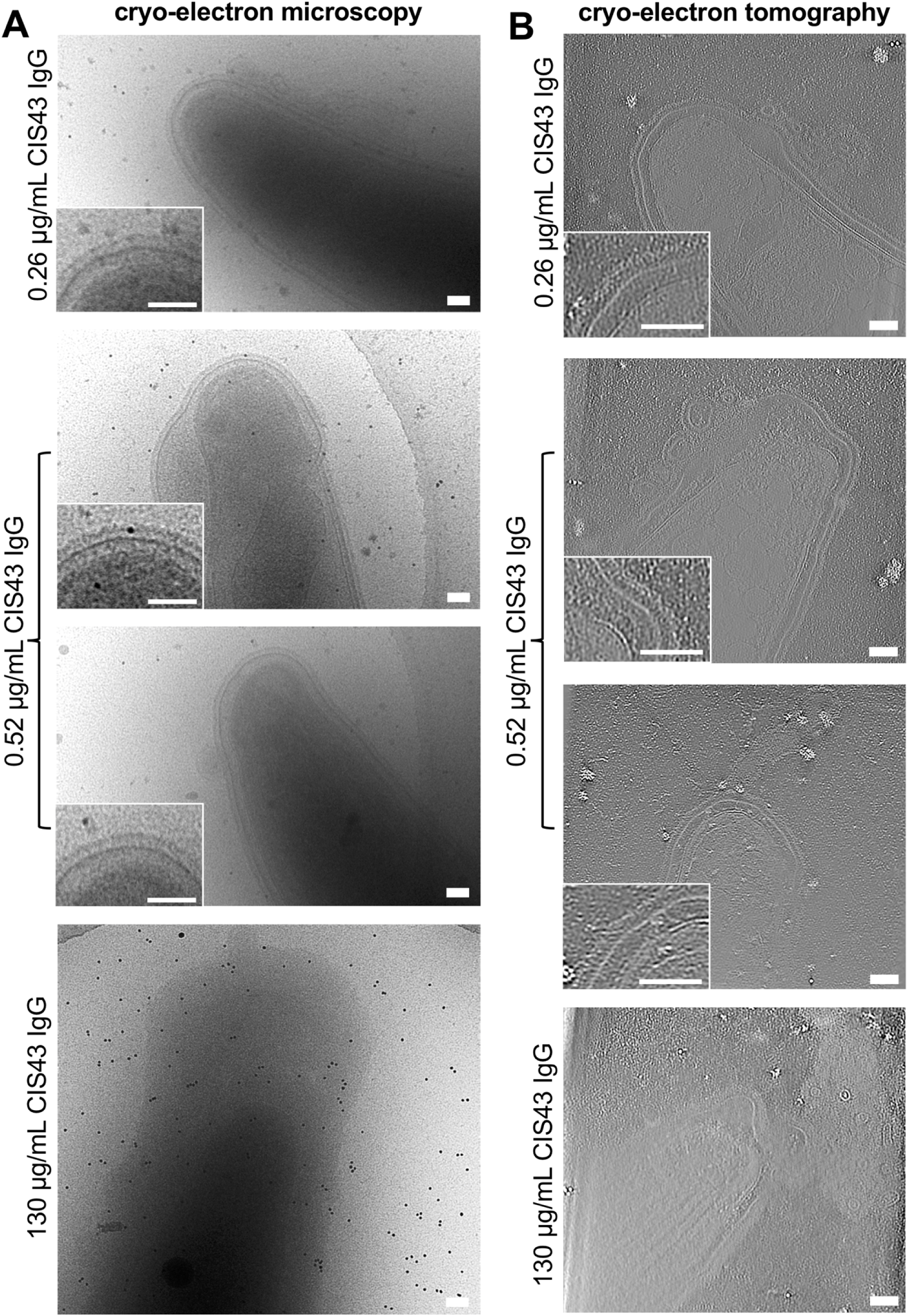
Titration of CIS43 IgG onto whole PfSPZ. At 0.26 µg/mL and 0.52 µg/mL, an extra layer of density is seen near the cell surface. While at higher concentrations a CSP shedding reaction results. **(A)** Cryo-EM imaging; insets show close-up of sporozoite surface after CIS43 IgG incubation. **(B)** Cryo-ET imaging showing central tomographic slices with inverted contrast; inset shows magnified surface. Additional examples are provided in Supplemental Figures S3 and S4. All scalebars 100 nm.

**Fig 3.**
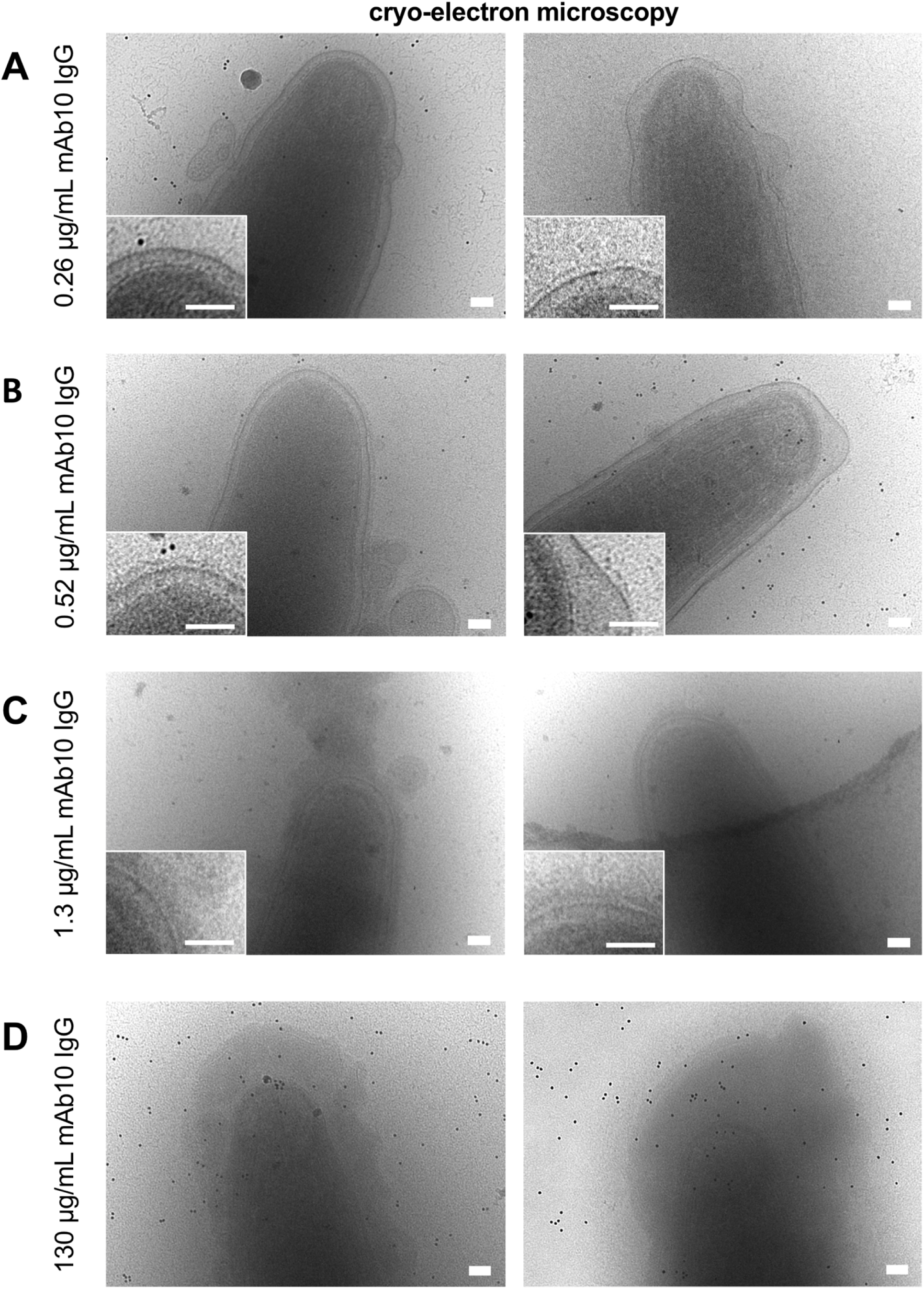
Titration of mAb10 IgG onto whole PfSPZ imaged by cryo-EM. At 0.26 µg/mL **(A) and** 0.52 µg/mL **(B)** an extra layer of density is seen near the cell surface. While at 1.3 µg/mL **(C)** a thick density layer as well as initiation of shedding is observed. **(D)** At 130 µg/mL extensive CSP shedding is observed. Insets show close-up of sporozoite surface after mAb10 IgG incubation. Additional views provided in Supplemental Figures S5, S6, and S7. All scalebars 100 nm.

With the incubation of CIS43 IgG at the lowest concentration (0.26 µg/mL), a thin, relatively uniform layer of density was observed along the surface of the PfSPZ (**Fig 2**). In cryo-EM projection images, this layer was positioned 27.3 ± 2.7 nm above the cell outer membrane (**S3 Fig**). When we double the concentration of CIS43 IgG (0.52 µg/mL), the layer thickened and extended slightly further to 29.4 ± 3.0 nm away from the surface (**Fig 2 and 4, S4 Fig**). Increasing CIS43 IgG concentration to 1.3 µg/mL during incubation, we found that the sample became too thick and too electron dense to image (data not shown), suggesting that the PfSPZ cells were blanketed in shed CIS43 IgG-bound CSP protein resulting from the CSP reaction. When we image PfSPZ that have incubated with 130 µg/mL of CIS43 IgG, a concentration comparable to what is anticipated towards the high end of antigen specific antibody concentrations in vivo [64], although we had difficulty finding cells with sufficient beam penetration to image, it was possible to obtain views of areas where a PfSPZ cell was immersed in a thick cloud of shed PfCSP and IgG, a distended membrane, and in some cases small membrane-bound vesicles (**Fig 2**). We note that there may be several stages leading up to CSP reaction, and that this reaction may be antibody concentration-dependent, antibody-specific, and environment-dependent. It is possible for the PfSPZ cell to separate from the density of IgG and PfCSP complex at high concentrations of central repeat-targeting cytotoxic antibodies, prior to the SPZ becoming unable to carry out further infection. A report from Aliprandini et al observed incubation with central-repeat binding antibodies will induce parasite death that is associated with a “thread-like” precipitate at the posterior end of GFP-expressing *P.yoelii* SPZ [54]. We speculate that a thick plume may start to accumulate around the cell at 1.3 µg/mL; and that at 130 µg/ml, the CSP coat begins to shed away from the cell body. Cryo-ET of the CIS43-PfSPZ samples across the antibody concentrations revealed similar features (**Fig 2B**). Given that cryo-EM could provide similar information as cryo-ET, moving forward, we used cryo-EM for the remaining antibody studies.

**Fig 4.**
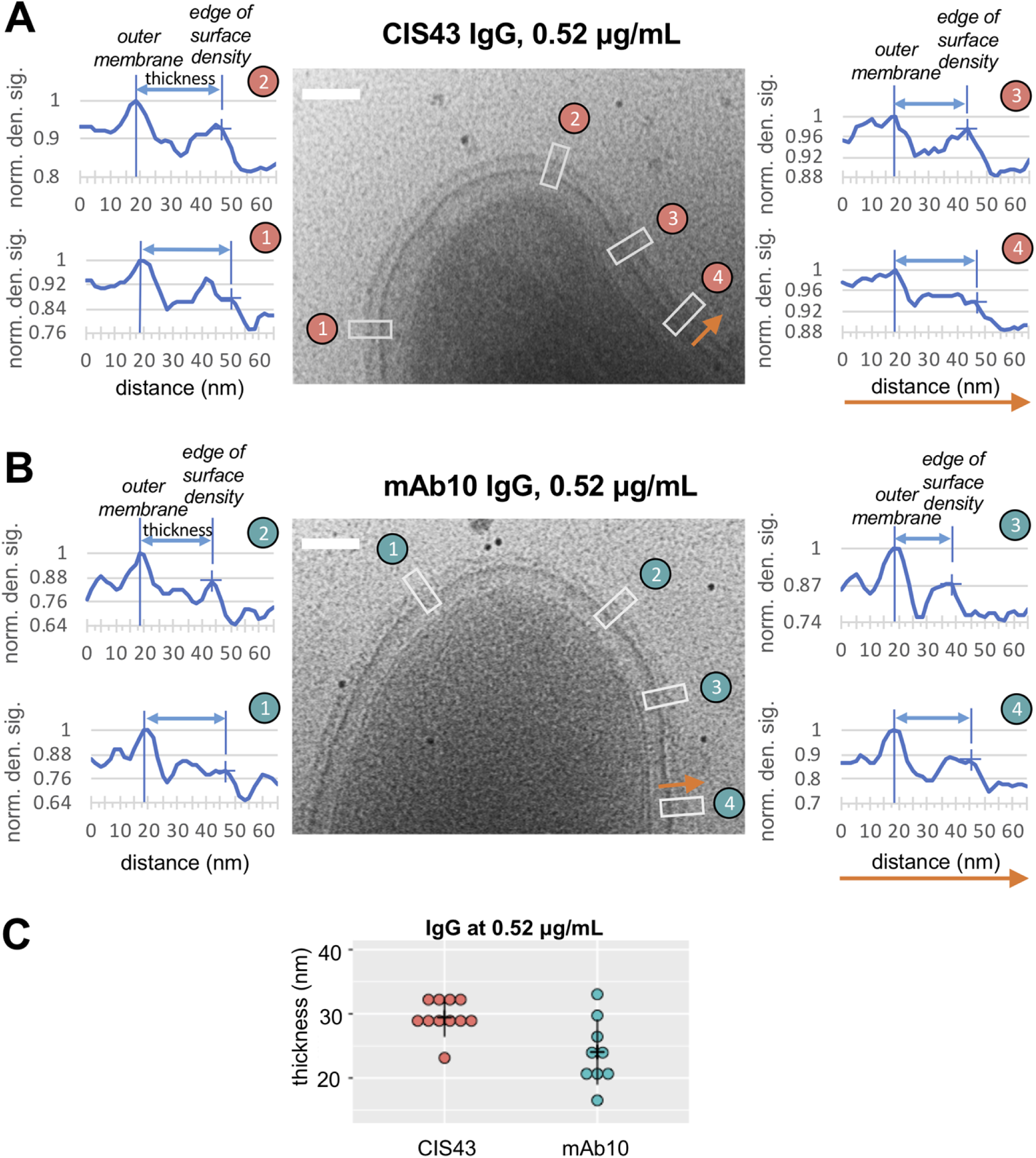
Analysis of surface density on PfSPZ following incubation with CIS43 or mAb10 IgG. Electron density greyscale plots across the distances of the boxed regions in cryo-EM micrographs of PfSPZ incubated with **(A)** 0.52 µg/mL CIS43 IgG (salmon plots) or **(B)** 0.52 µg/mL mAb10 IgG (teal plots). The orange arrows indicate direction of measurement on x-axis. The distance of the surface thickness is measured between the PfSPZ outer membrane and the edge of the surface density. **(C)** Dot plot shows surface layer thickness measurements from multiple PfSPZ at 0.52 µg/mL of CIS43 or mAb10 IgG. Additional measurements are provided in Supplemental Figures S4 and S7. Scalebars 100 nm.

mAb10 IgG, which targets the NANP central repeats, exhibited slightly different trends in binding compared to CIS43 IgG. When incubating at the lowest concentration of 0.26 µg/mL, instead of a uniform density surrounding the cell, mAb10 produced a heterogeneous population of cells where some cells exhibited a patchy layer of density coating the surface, while others remained apparently bare of CSP-bound IgG (**Fig 3, S5 and S6 Fig**). At 0.26 µg/mL on the cells with patches of surface density, the layer extended to 21.5 ± 5.2 nm above the outer cell (**S6 Fig**), which is several nm shorter than the layer observed in CIS43, consistent with mAb10 binding to a more membrane-proximal positioning of the central repeats than CIS43, which preferentially binds first to the Junction epitope over to the central repeats [36]. When mAb10 was present at 0.52 µg/mL, the cells exhibited a uniform density layer on the surface with an average thickness of 24.0 ± 5.1 nm (**Fig 3 and 4, S7 Fig**). As mAb10 IgG concentration was further increased to 1.3 µg/mL, the cells showed signs of undergoing CSP shedding reactions, where a trail of density was observed from some of the SPZ ends (**Fig 3**). In contrast to what we observed with CIS43 IgG incubation at this concentration, with 1.3 µg/mL concentration of mAb10 bound, the electron beam was still able to penetrate the proteinaceous density around the cell, and the cell body was still discernable inside (**Fig 3**). At 130 µg/mL mAb10 IgG incubation concentration, the sporozoites were showed large, electron-dense clouds of IgG bound to PfCSP sloughing off in a similar fashion to CIS43 incubation at this same concentration (**Fig 3**).

#### Monoclonal antibody mAb15 binds at low levels at a membrane-proximal CTD position

We also examined a PfCSP CTD antibody, mAb15 IgG, which was originally isolated from a human subject who was immunized with the PfSPZ vaccine [36]. We observed no apparent bound density on the surface of the sporozoite at 0.52 or 1.3 μg/mL concentrations of mAb15 (**S8 Fig**). The CSP α-thrombospondin type-1 repeat (αTSR) is anticipated to be membrane proximal and located within the CTD. While the 67-residue αTSR is too small to clearly resolve by cryo-EM, the structure of the isolated, recombinant domain has been previously determined by X-ray crystallography [21]; despite the potential added density of mAb15 at these concentrations, there is no detectable density on the surface of the cells. However, at 130 μg/mL, an additional density layer of relatively uniform thickness was observed adjacent to the membrane surface on most sporozoites; this layer is evident as a shoulder in a density plot, thickening the SPZ outer membrane by ∼10-15 nm compared to what is observed with no antibody present (**Fig 5, S9 Fig**). This observation is consistent with mAb15 binding close to the SPZ surface where the CTD is located. With mAb15 IgG, despite the high antibody concentration and a uniform coating of the SPZ surface by bound antibody, no apparent large-scale CSP shedding reaction was observed.

**Fig 5.**
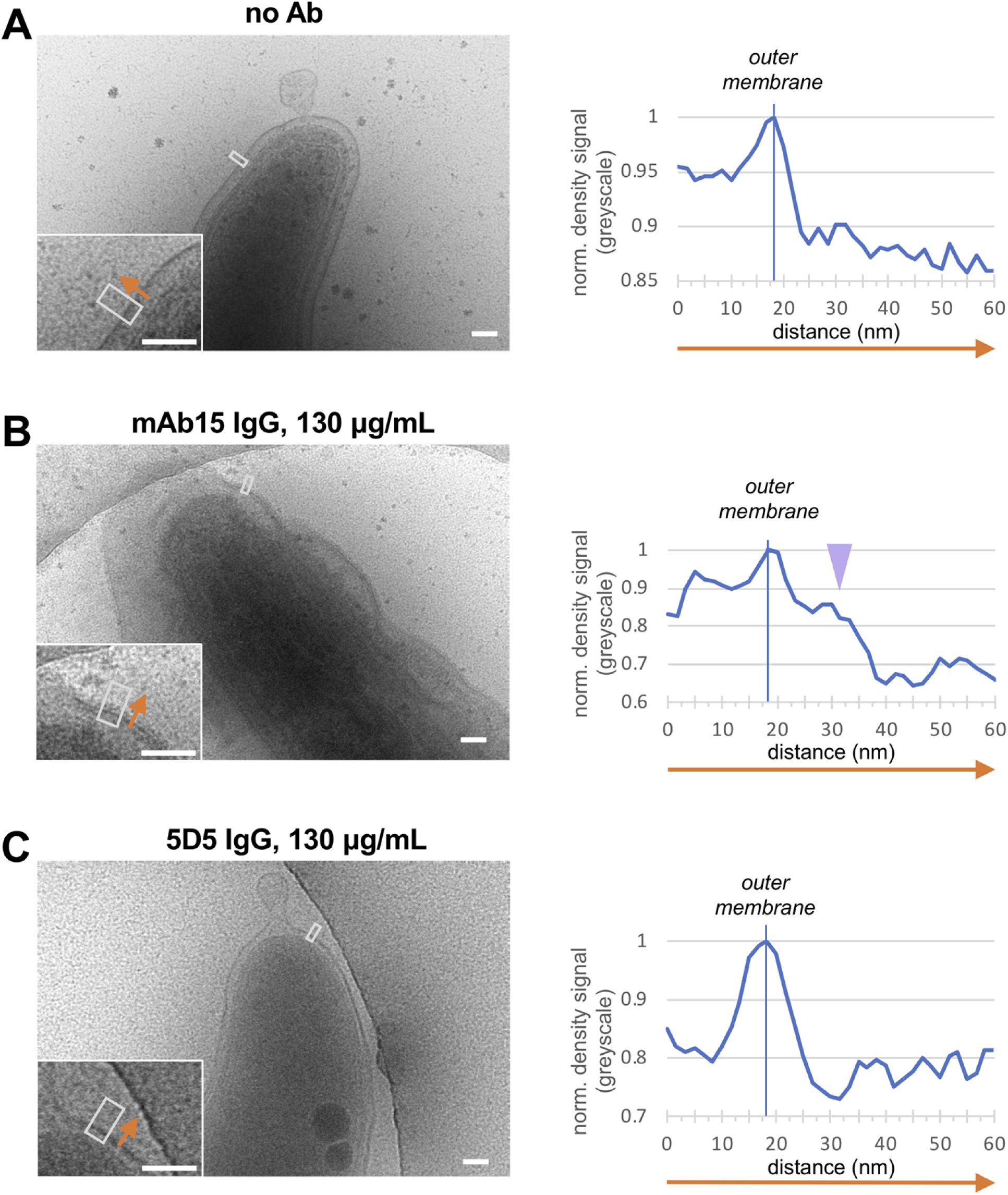
Imaging PfSPZ with NTD and CTD-targeting antibodies. **(A)** Antibody naïve PfSPZ, **(B)** following incubation with 130 µg/mL mAb15 IgG, **(C)** following incubation with 130 µg/mL 5D5 IgG. Orange arrow indicates the direction of density measurement on the cryo-EM micrograph. **(B)** In PfSPZ incubated with 130 µg/mL mAb15 IgG, extra density in the form of a diffuse layer on the surface is seen compared to the antibody-naïve case; the lilac arrowhead points to the shoulder that extends out from the PfSPZ surface and thickens the outer membrane as a result of mAb15 IgG binding. **(C)** In PfSPZ incubated with 130 µg/mL 5D5 IgG, the surface does not appear to differ relative to the antibody-naïve control. Scalebars 100 nm. Additional examples provided in Supplemental Figure S9.

#### Monoclonal antibody 5D5 against the NTD does not bind appreciably to whole radiation-attenuated PfSPZ

5D5 is an NTD-binding antibody that was isolated following murine immunization using *P. berghei* transgenic parasites expressing a chimeric CSP that contains the N-terminal region of *P. falciparum* CSP [31]. Its epitope has been traced using ELISA with a panel of CSP-spanning peptides and confirmed in a crystal structure of 5D5 FAb bound to a peptide that is found N-terminal to the Junction region in the PfCSP NTD [31, 35]. We observed that in the presence of 130 μg/mL of 5D5, the surface of the sporozoites appeared essentially identical to antibody-naïve sporozoites (**Fig 5**, **S9 Fig**), consistent with the finding by flow cytometry that 5D5 binds at negligible levels to PfCSP on live *P. falciparum* sporozoites, despite the antibody’s high binding affinity to its cognate peptide sequence in isolation [35].

#### Repeat antibody FAb’s coat the sporozoite surface without inducing major shedding or the CSP reaction

From the continuity of density layers seen in the presence of antibodies, we can infer that PfCSP is densely arrayed on the sporozoite surface. With central-repeat antibodies, the possibility exists for bivalent intra- and inter-CSP cross-linking of IgG, which then would be multiplied across the PfSPZ surface. In order to examine the effect of antibody bivalency on sporozoites and the CSP reaction, we incubated PfSPZ with CIS43 and mAb10 FAb fragments. At 130 μg/mL incubation, the cell surfaces exhibited dense coats of relatively even thickness, in stark contrast to the dramatic plumes of shed density seen for IgG bound at equivalent concentrations in the CSP reaction **(S10 Fig**). For CIS43 FAb, the average thickness of this density layer, measured from the cell outer membrane to the edge of the surface density, was 30.9 ± 4.2 nm (**Fig 6, S11 Fig**). For mAb10 FAb, the thickness of the density layer on the cell surface was slightly greater, averaging 34.9 ± 3.5 nm (**Fig 6, S12 Fig**). These dimensions are measurably thicker than the layers seen for the intact IgG, suggesting that monovalent FAb’s likely were able to more completely saturate the repeats and extend CSP-FAb density outward from the surface.

**Fig 6.**
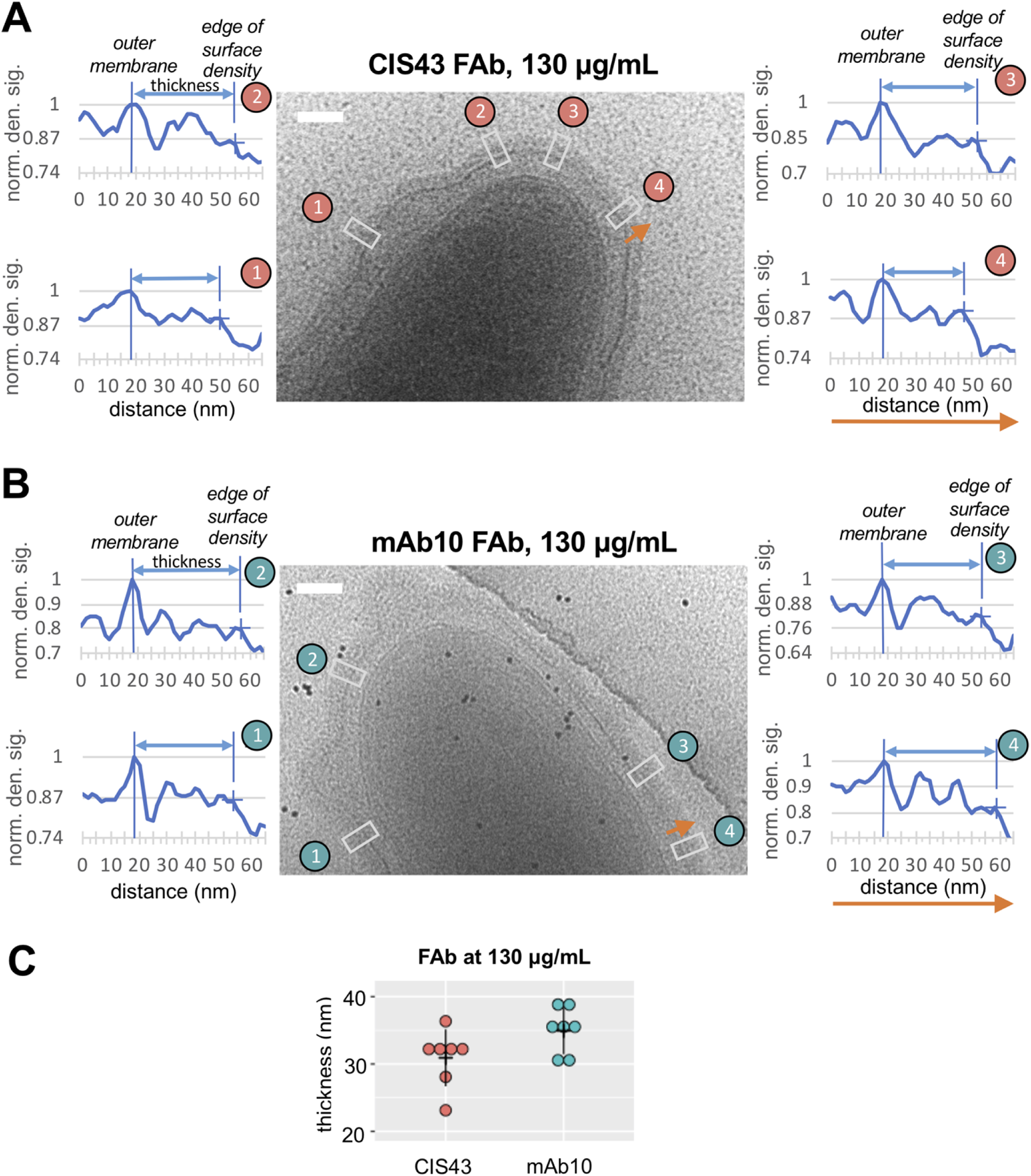
Role of anti-CSP repeat antibody bivalency on CSP binding and shedding. Electron density greyscale plots across the distances of the boxed regions in cryo-EM micrographs of PfSPZ incubated with **(A)** 130 µg/mL CIS43 FAb (salmon plots) or **(B)** 130 µg/mL mAb10 FAb (teal plots). The orange arrows indicate direction of density measurements on the x-axis. The surface layer thickness is measured between the SPZ outer membrane and the edge of the surface density. **(C)** Dot plot shows surface layer thickness measurements (nm) from multiple SPZ at 130 µg/mL of CIS43 or mAb10 FAb. Additional measurements and views provided in Supplemental Figures S10, S11, and S12. Scalebars 100 nm.

For both FAbs, there was no observable shedding in the form of a large-scale CSP reaction, indicating that the CSP reaction relies on the bivalency of repeat antibody IgG’s, due to their ability to crosslink and network PfCSP together across the PfSPZ surface. We do note that differences in FAb-dependent effects resulting from binding of CIS43 and mAb10 FAb were observed, however. With the CIS43 FAb binding, a subpopulation of PfSPZ were observed where the cell membrane appears to inflate, expanding the inner membrane complex compartment substantially, while retaining an intact outer membrane layer coated evenly by the FAb-bound CSP density (**S10A Fig**). With the mAb10 FAb, by contrast, a number of sporozoites exhibited small vesicles with relatively uniform layers of FAb-bound PfCSP surrounding the cell. These vesicles appear to possess intact membrane bilayers that have budded or blebbed from the cell outer membrane. Indeed, more commonly seen for mAb10 FAb-bound sporozoites are regions where the outer membrane is buckling outward, perhaps in early stages of forming a vesicle (**S10B Fig**).

A previously study [37] using single particle cryo-EM to study FAb and IgG binding to a truncated recombinant PfCSP reported that certain repeat antibody FAb’s can adopt a highly ordered structure consisting of multiple copies of FAb stacked into a spiral structure. The reported FAb+PfCSP assemblies were measured to have a width of approximately 8 nm [37]. Because several of our whole PfSPZ bound with central repeat-binding FAb’s (CIS43 and mAb10) images show density on the surface that could be interpreted as columns of density extending perpendicular to the sporozoite surface (**S10 Fig**), we sought to compare these *in situ* features with the assemblies described in the previous report [37]. Particularly for mAb10 FAb, the columnar features were seen to be relatively regular in side-by-side spacing and in overall dimensions (**Fig 7A**). Plotting the density parallel to the surface of the cell in the boxed areas, we observed repeating peaks that correspond to relatively regular lateral positioning of the columnar features (**Fig 7**). The width of each peak, measured as the width (nm) at the half maximum height of the density peak, was found to be on average 7.6 ± 1.7 nm for mAb10 FAb, consistent with the dimensions observed in the single particle FAb-recombinant CSP assemblies [37] (**Fig 7B**). CIS43 FAb incubation with SPZ also showed similar features in certain regions, with peak widths of 7.7 ± 2.5 nm (**Fig 7C**), however these were somewhat less distinct than observed for mAb10. These observations suggest that FAbs can form stacked spirals with high occupancy levels of the PfCSP repeats on surface of SPZ. No such features were observed, however, with intact bivalent IgG, most likely due to their ability to bind repeat epitopes both within a PfCSP copy and to span adjacent PfCSP, giving rise to a less ordered antibody-PfCSP complex on the SPZ surface that is not permissive to high levels of regular FAb-FAb stacking.

**Fig 7.**
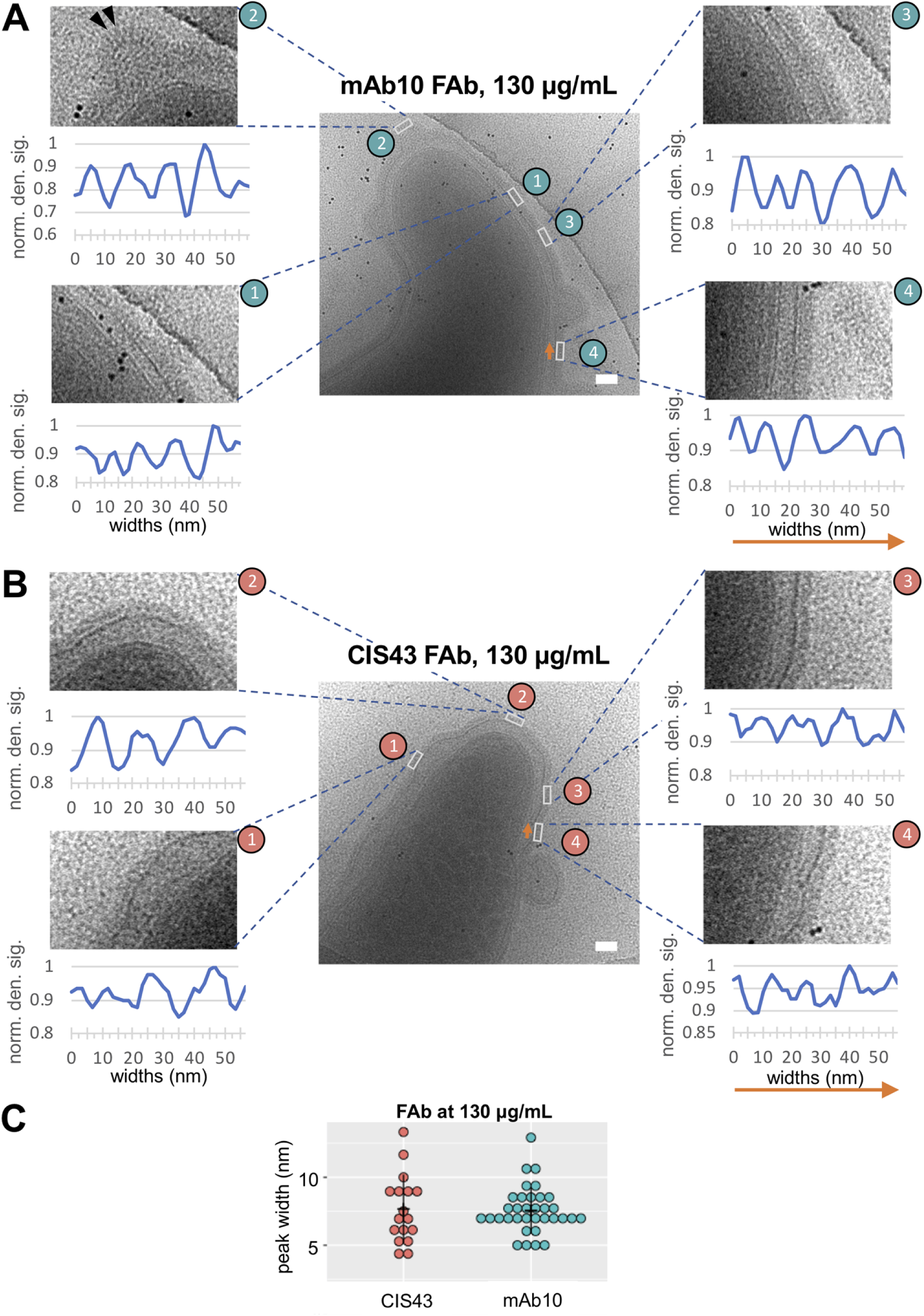
Formation of higher order repeat-antibody Fab-CSP features on PfSPZ following incubation with mAb10 FAb or CIS43 FAb. Electron density greyscale plots across the distances of the boxed regions in micrographs of PfSPZ incubated with **(A)** 130 µg/mL mAb10 FAb (teal plots) or **(B)** 130 µg/mL CIS43 FAb (salmon plots). The orange arrows indicate the direction of measurement along the PfSPZ surface, parallel to the cell surface. The plots reveal a regularly spaced pattern of density peaks. Peak widths measured at half-max peak height (black arrowheads) in the micrographs. **(C)** Dot plot shows the peak widths (nm) of the surface column features from measurements along the SPZ surface. Scalebars 100 nm.

## DISCUSSION

Developing a basic understanding of how antibodies recognize CSP on *Plasmodium* sporozoites and which epitopes are accessible in the antigen’s native context provides valuable information that can guide anti-malarial vaccine development. By examining the binding effects of CIS43 and mAb10 IgG’s, we estimate that the Junction of PfCSP is located farther away from the outer membrane than the central repeats. The extra densities at the lowest concentrations of CIS43 IgG [36, 65] extend ∼27-29 nm above the PfSPZ outer membrane (**Fig 4, S3 Fig**), which we interpret to reflect CIS43 initially binding to its primary high-affinity Junction epitope preferentially over the central repeat secondary epitopes. By contrast, mAb10 exhibits a single binding mode to the central repeats [36, 65]. Based upon our analysis of mAb10 binding, the furthest edge of the central repeat region thus appears to be ∼5-6 nm shorter and closer to the outer membrane than the Junction (**Fig 4, S6 Fig**). Since the Junction appears to be accessible on whole SPZ, inclusion of this motif on CSP based immunogens is likely to aid in eliciting protective antibodies.

Both of the repeat-binding antibody IgG’s are shown to induce a CSP reaction where an IgG-CSP precipitate is shed off the cell surface. We observed this CSP reaction is dependent on two conditions: exceeding a threshold concentration of a central-repeat binding antibody and the bivalency of the antibody. We find that 1.3 µg/ml concentration of CIS43 and mAb10 is sufficient to begin to induce the CSP reaction on whole PfSPZ. CIS43 IgG at 1.3 µg/mL results in a strong CSP reaction exhibiting broad, diffuse plumes where the shed density completely envelopes the cell and the sample becomes too thick to image by cryo-EM and cryo-ET. By comparison, incubation with 1.3 µg/ml mAb10 IgG results in a thinner, more compact plume that is shed that allows the beam to penetrate the sample (**Fig 3**). The dual binding modality of CIS43 IgG compared to the single mode binding of mAb10 IgG may contribute to the differences we observe in the CSP reaction phenotypes. These observations by cryo-EM are consistent with a recent study by Guiterre-Botero et al. that used fluorescence microscopy to show that CIS43 induces a CSP reaction that yields a condensed bolus of shed CSP with a “short-round phenotype” according to the authors, while mAb10 produces an elongated coat that retains the shape of the SPZ cell [65]. The concentrations of antibody that we observe to induce shedding are within the range of CSP-specific antibodies that have been observed in serum following vaccination, suggesting it is possible to elicit sufficient concentrations of antibodies that can disrupt the CSP coat and its essential functions that the sporozoite relies upon for infection [54, 64].

CSP reaction shedding was also demonstrated to be dependent upon IgG bivalency. When FAb fragments of CIS43 and mAb10 were incubated, even at very high concentrations, shedding on PfSPZ was not observed (**Fig 6**). Instead, a relatively uniform coat of density resulting from FAb condensing on PfSPZ produced relatively orderly features (**Fig 7**) that are consistent with previously described columns of stacked FAb domains bound to NANP repeats in recombinant PfCSP, where the PfCSP peptide was organized in a spiral towards the center of the column of density [39, 66]. We observe that mAb10 FAb produced more defined CSP ordering on the PfSPZ surface than CIS43 FAb (**Fig 7**), which may reflect CIS43’s dual binding modality [36, 65]. While CSP reaction shedding was not observed here with FAbs, it did appear that high levels of FAb binding to PfCSP on PfSPZ could on some cells produce a degree of outer membrane buckling and vesiculation, specifically with mAb10 FAb (**S10B Fig**). While certain repeat-targeting IgG’s may mirror the stacking spiral organization seen with FAb’s along a recombinant PfCSP peptide [37, 39, 66], on the surface of sporozoites where PfCSP protein is present at high densities, inter-CSP crosslinking by bivalent IgG most likely competes with any intra-CSP binding, and will induce far greater perturbation of the PfSPZ surface resulting in a shedding CSP reaction compared with the same concentrations of FAb. At those same concentrations of FAb, we find no CSP reaction and supports that the bivalency of IgG contributes to a CSP reaction. The thinner surface density layer seen with CIS43 FAb (∼31 nm) versus mAb10 FAb (∼35 nm) may reflect the lower binding site saturation observed by Aguirre-Botero et al for CIS43 antibody [65].

The CSP reaction shedding is not expected of antibodies that have a single binding motif present such as with mAb15. While we observed additional density due to mAb15 binding close to the sporozoite outer membrane (**Fig 5B**), and while it may be possible for one IgG to crosslink an adjacent CTD on a neighboring PfCSP, higher order networks of crosslinked PfCSP and antibody would not be able to form for such CTD-targeting antibodies. Such higher order crosslinking appears to be necessary to induce the CSP reaction. We primarily observed a layer of density we attribute to mAb15 bound to the CTD at the highest concentrations tested, 130 µg/mL. By contrast, Kisalu et al observed that mAb15 could bind to soluble, recombinant PfCSP at far lower concentrations, e.g. 5. µg/mL, as detected by ELISA [36]. We infer that the apparently intrinsically disordered and densely-packed PfCSP on the PfSPZ surface likely presents a steric barrier for CTD-targeting IgG’s to access the membrane proximal epitope at lower concentrations [30, 49]. High concentrations are required to enable sufficient amounts of mAb15 to overcome this barrier so as to be observable by cryo-EM.

Among the various epitopes on PfCSP, the NTD has been a somewhat enigmatic target, with different studies reporting disparate results. Antibody 5D5, generated from mouse immunizations with full-length, recombinant PfCSP [31], has been shown to recognize a linear epitope spanning residues 82-91 in the NTD of PfCSP based upon previous studies measuring binding to CSP-derived peptide fragments [34]. This sequence is located N-terminal to the cleavage site between Lys95 and Gln96 residues, a proteolytic processing site purported to be necessary for the later hepatocyte invasion stage of malaria infection [30, 31, 35]. Our examination of sporozoites following incubation with 5D5 at high concentrations revealed no observable density that could be attributed to the NTD-targeting antibody (**Fig 5C**). This is observation is consistent with previous reports that show the vast majority of freshly dissected sporozoites do not bind fluorescently-labeled 5D5 antibody [35]. Recently, Dacon et al reported analysis of sporozoites where 5D5 was shown to only bind to uncleaved forms of PfCSP from *Plasmodium falciparum* sporozoite lysate [34]. Our observations indicate that the radiation-attenuated PfSPZ in our study predominantly display a cleaved form of PfCSP that lacks the 5D5 epitope.

The studies we report here highlight how PfCSP and its epitopes are presented on whole sporozoites and elucidate the effect of PfSPZ-antibody interactions on the integrity of the parasite’s CSP coat and outer membrane layer. Studying CSP in its native environment is of significant interest for optimizing immunogen design, since whole attenuated sporozoite immunization has been shown to provide long-lasting protection against malaria disease [7–12]. It is worth noting the current the RTS,S/AS01 (Mosquirix) vaccine includes a truncated version of PfCSP, leaving out the important Junctional region as well as the minor repeats that can generate strong protective antibodies against those epitope regions [36, 39, 47]. Recent studies suggest that administration of Junction-targeting antibodies together along with immunization of R21, another truncated PfCSP-based vaccine that elicits antibodies against the repeat region and CTD of CSP, provides greater protection against SPZ challenges than vaccine or Junction-targeting antibody alone [67, 68]. Additionally, it has been reported that inclusion of the Junction and minor repeats on immunogens induce protective antibodies *in vivo* [69]. In future vaccine designs, it may be beneficial to include the Junction region along with the central repeats in order to elicit responses against both of these key epitopes to provide broad and lasting protection.

In summary, the results of this study shed light on the accessibility of native PfCSP epitopes on radiation-attenuated *Plasmodium falciparum* and the effect the antibodies have on the CSP coat and sporozoite morphology, which can inform future malaria vaccine design and development.

## METHODS AND MATERIALS

### Preparation of sporozoites on cryo-EM grids

Live attenuated *P. falciparum* sporozoites (PfSPZ) were purchased from Sanaria (150,000 per 20 µL vial) and washed by adding 50 µL 1% of human serum albumin (HSA) in PBS, followed by centrifuging to pellet SPZ (13,000 x g, 3 minutes), carefully removing top 50 µL, and resuspending the pellet in the remaining liquid. Washed SPZ are incubated with either IgG or FAb at room temperature for 10 minutes, then allowed to sit on carbon-coated Quantifoil grids (3.5/1, 300 mesh, copper) with 10 nm gold beads (Nanoprobes) for additional 10 minutes before plunge freezing in Vitrobot (100% humidity, 4 °C, 11 seconds blot time).

### Collection and processing of cryo-electron microscopy and cryo-electron tomography

Cryo-electron micrographs were collected on a T12 Spirit with 30 e^-^/A^2^ total dose and a - 12 µm defocus (4.13 Å/pixel). A high defocus was necessary to image these large PfSPZ cells with a 0.5 µm cross-sectional thickness. Micrographs of samples are depicted on an electron density greyscale signal. The boxed regions in the micrographs are presented as electron density grey scale plots using ImageJ tools, with the maximum density signal value normalized to a value of 1, and all other values scaled relatively. Cryo-electron tilt series frames were collected on Krios with K2 camera and a 20 eV energy filter, 50 e^-^/A^2^ total dose and a -10 µm defocus, ±48°, 3° step (3.25 Å/pixel). Frames were aligned with MotionCorr2 and the aligned tilted series were reconstructed with IMOD.

### Antibody/FAb expression and production

Genes encoding antibody heavy and light chains were cotransfected in HEK293E (RRID:CVCL_HF20) cells at 1×10^6^ cells/mL using a total of 500 μg DNA per liter of culture (1:1 ratio of HC-to-LC DNA) using 293 Free Transfection Reagent (Novagen). Six days following transfection, cells were centrifuged at 4,500 rpm for 20 minutes. Supernatant was filter sterilized 0.22-µm filter and incubated with Protein A (Thermo Fisher) resin overnight at 4°C on a shaker. The resin was separate from the supernatant and washed with five column volumes of 1x PBS and eluted with IgG Elution buffer (Thermo Fisher). Tris (1 M) was added immediately to neutralize the pH to ∼7.4, subsequently buffer exchanged into a 20 mM HEPES/HCl pH 7.5, 150 mM NaCl buffer using a 30kDa concentrator (Millipore). The IgG’s were run on size-exclusion chromatography using a HiLoad 16/600 Superdex 200 pg (GE) column using the same buffer.

To produce antigen-binding fragments (FAb’s), purified IgG’s were digested with LysC (New England Biolabs) overnight at 37 °C (1 μg for 10 mg of IgG). To remove LysC and any undigested IgG, the FAb’s were purified using a HiLoad 16/600 Superdex 200 size exclusion column (GE) on an AKTApure system (GE) into a buffer containing 20 mM HEPES/HCl pH 7.5, 150 mM NaCl.

## DATA AVAILABILITY

All data and micrographs are available upon request.

## ACKNOWLEDGEMENTS

We thank Dr. Azza Idris (Massachusetts Institute of Technology) for helpful comments and Dr. Robert Seder (NIH) for generously providing some of the sporozoites used in this study. This work was supported by grants INV-084290, INV-010646, and INV-043758 from the Gates Foundation. We thank the University of Washington Arnold and Mabel Beckman Cryo-EM Center for data collection time and support.

## AUTHOR CONTRIBUTION

Nancy Hom

Contribution: Conceptualization, Data curation, Formal analysis, Investigation, Methodology, Visualization, Writing original draft, Writing review and editing

Connor Weidle

Contribution: Methodology, Writing review and editing

Marie Pancera

Contribution: Conceptualization, Funding acquisition, Project administration, Resources, Supervision, Writing review and editing

Kelly K. Lee

Contribution: Conceptualization, Data curation, Formal analysis, Funding acquisition, Project administration, Resources, Supervision, Visualization, Writing original draft, Writing review and editing

